# Systemic post-translational control of bacterial metabolism regulates adaptation in dynamic environments

**DOI:** 10.1101/180646

**Authors:** Elizabeth Brunk, Roger L. Chang, Jing Xia, Hooman Hefzi, James T. Yurkovich, Donghyuk Kim, Evan Buckmiller, Harris Wang, Chen Yang, Bernhard O. Palsson, George M. Church, Nathan E. Lewis

## Abstract

Across all domains of life, elaborate control mechanisms regulate proteins, pathways, and cell phenotypes as organisms adapt to ever-changing environments. Post-translational modifications (PTMs) allow cells to rapidly and reversibly regulate molecular pathways, but it remains unclear how individual PTMs regulate fitness. Here, we studied >130 PTM sites in *Escherichia coli* to unravel how PTMs regulate cell metabolism and fitness in response to environmental changes, such as the glucose-acetate diauxie. Using a new metabolic modeling approach, we found a significant fraction of post-translationally modified enzymes are predicted to control shifts in pathway usage following evolutionarily-important environmental changes. Genetic screens using Multiplex Automated Genome Engineering confirmed that these PTMs impact cellular fitness, especially under dynamically changing environments. Finally, mechanisms of how individual PTMs impact protein function were detailed using molecular dynamics simulations and enzyme assays for enolase, transaldolase, and serine hydroxymethyltransferase. Thus, by integrating whole-cell data and pathway modeling with detailed biochemical analysis, we unraveled how individual PTMs regulate enzymes, pathways, and phenotypes to adapt to sudden environmental changes.

## Introduction

Organisms have evolved elaborate control mechanisms to regulate cell physiology and help cells survive and compete as environments frequently and suddenly change(López-Maury et al., 2008; Savageau, 1998). For example, bacteria that are consumed by an animal will be exposed to several rapid nutritional shifts (e.g., oxygen, sugar, and amino acid levels) as they traverse through the gastrointestinal track. Familiar changes that are sustained for longer periods of time will invoke transcriptional regulatory mechanisms to gradually adapt a cell to an environment. However, these mechanisms are costly in time and resources, and are inadequate to respond to sudden and transient changes from environmental perturbations or expression noise in the cell. To cope with immediate environmental changes in metabolism, small metabolites directly regulate enzymes through allosteric or competitive mechanisms.

Post-translational modifications (PTMs) are of particular interest since they can rapidly regulate cell pathways, but their effect is also more sustained than regulation through metabolite feedback(Kochanowski et al., 2015). This is particularly important to adapt to nutritional fluctuations in the microenvironment or transient variations in enzyme abundance in a single cell (Labhsetwar et al., 2013). Despite their potential importance, many PTMs remain poorly characterized, and their regulatory roles remain unstudied. Indeed, only about a half dozen metabolic enzymes in *E. coli* are specifically known to be regulated by PTMs to control metabolic flux (Pisithkul et al., 2015). Furthermore, in prokaryotes, it is commonly assumed that phosphorylation is primarily relevant to two-component signaling (Pearlman et al., 2011). Recent studies, however, challenged these perceptions by elucidating the widespread nature of PTMs in bacteria, which suggest that many more bacterial proteins are regulated by acetylation or phosphorylation(Cain et al., 2014). While some PTMs could be spurious and low-stoichiometry chemical modifications(Weinert et al., 2013), there remains compelling evidence that many PTMs could regulate bacterial metabolism(Chubukov et al., 2014; Hansen et al., 2013; Kochanowski et al., 2015). However, it has been difficult to unravel the physiological roles of the PTMs in a high-throughput manner. Thus, it is unclear (i) when PTMs are used to regulate metabolic enzymes, (ii) how the PTMs impact downstream pathways and regulate physiology, and (iii) how they exert their function on individual enzymes.

Here we demonstrate when and how many PTMs are deployed to control metabolism through the detailed analysis of >100 PTM sites in *E. coli*. We do this using several techniques including genome-scale metabolic modeling, genome editing using Multiplexed Automated Genome Engineering (MAGE), all-atom molecular dynamics simulations, and *in vitro* biochemical assays. Through this we demonstrate that: (i) PTMs occur on specific enzymes in bacterial metabolism that regulate important branch points; (ii) PTM-based regulation controls metabolic flux to enhance cellular fitness in dynamic environments; and (iii) PTMs alter the molecular properties of these proteins to modulate their activities. We further provide detailed analysis of PTMs on three specific proteins to highlight novel insights into how these proteins are regulated. Our findings further demonstrate how specific PTMs in bacterial metabolism can influence the fitness of an entire organism by facilitating adaptation to ever-changing environmental conditions.

## Results

### Metabolic regulation can be predicted in genome-scale models

When the nutritional environment changes, cells change their metabolic pathway usage accordingly(Chubukov et al., 2014) (Fig. S1-S2). Steady-state metabolic flux for all reactions can be predicted under diverse environments using constraint-based methods on genome-scale models (Lewis et al., 2012). The steady-state assumption of these methods, such as Flux Balance Analysis, specifically states that metabolite concentrations do not change. Therefore, it has been more difficult to use such approaches to study how the cell responds to sudden changes in the cellular microenvironment, which would require sudden transient regulation of specific enzymes. Here we overcome this challenge with a new algorithm, called Regulated Metabolic Branch Analysis (RuMBA), which predicts how cells regulate enzymes to optimally respond to transient metabolic perturbations.

The RuMBA algorithm specifically identifies enzymes that must be regulated to ensure optimal cell fitness immediately following a sudden change in its microenvironment. This is accomplished as follows. First, thousands of candidate flux states for all reactions are computed with Monte Carlo sampling, while enforcing near-optimal growth (Schellenberger and Palsson, 2009). This is done for the metabolic states before and after the perturbation. Second, these predicted flux states are subsequently compared (see Methods and Fig. S3). Specifically, each metabolite is queried to see if it is the center of a flux split (e.g., isocitrate in Fig. 1a). For each flux split, an empirical p-value is computed, testing if each reaction in the metabolic branch point significantly shifts its flux towards or away from the reaction. Thus, my taking a metabolite-centric view of Monte Carlo sampling data, RuMBA identifies enzymes that must be suppressed or activated to rapidly force flux from one branch to another to ensure optimal cell fitness immediately following the perturbation.

**Fig. 1.**
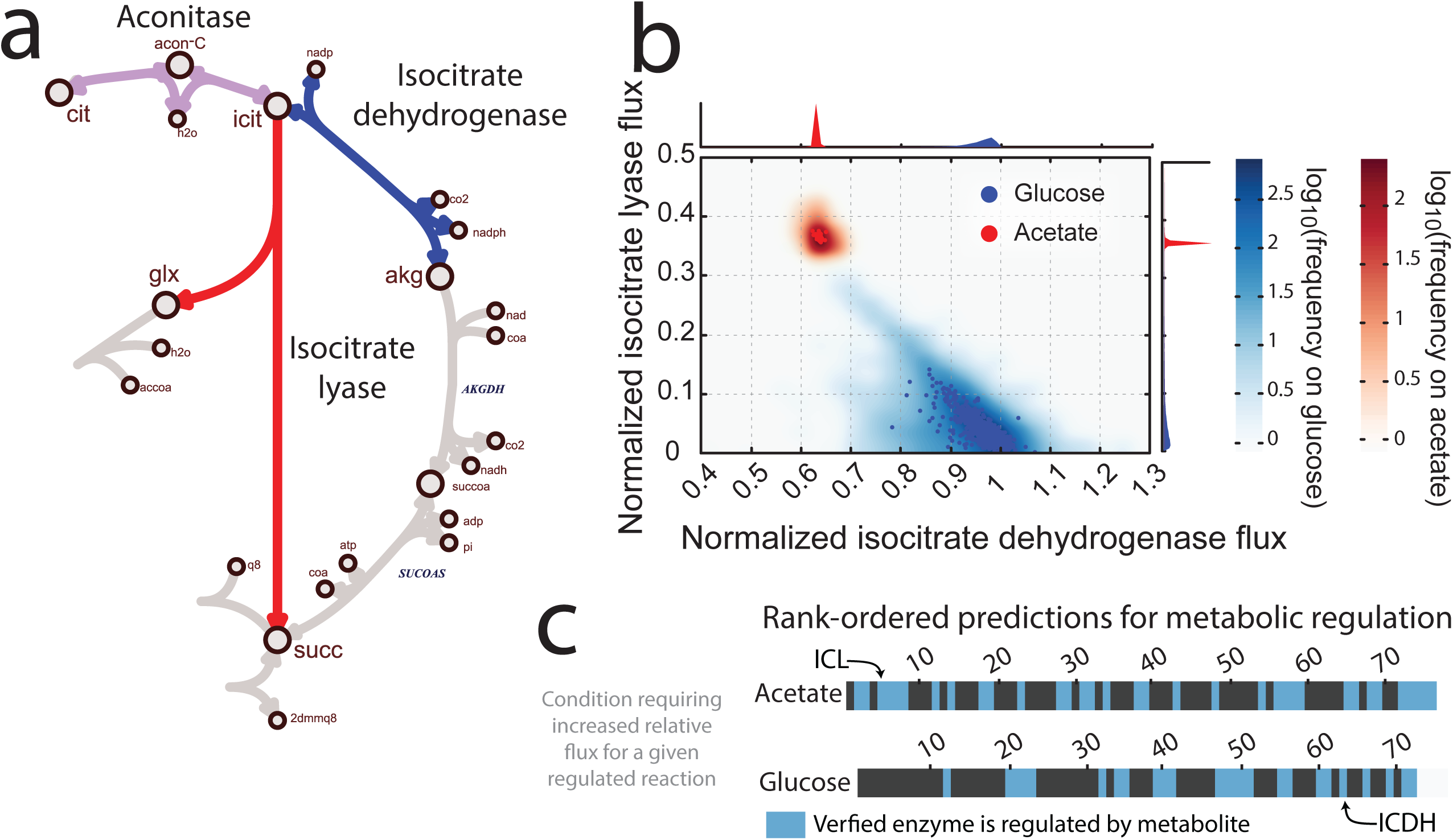
Metabolically-regulated enzymes can be predicted *in silico*. As changes in nutrient availability or enzyme copy number in cells occur, metabolic regulation can help rebalance metabolism while the transcriptional programs are deployed and fine-tuned. **(a)** This often occurs at branch-points in metabolism, such as the branch point at isocitrate, which divides flux between the TCA cycle and glyoxylate shunt during the glucose-acetate diauxie. **(b)** Randomly-sampled *in silico* flux distributions for growth on acetate and glucose show that isocitrate dehydrogenase is used for growth on glucose minimal media, while flux is diverted to isocitrate lyase when the cell is metabolizing acetate. **(c)** The RuMBA algorithm identifies a rank-ordered list of reactions and their associated enzymes that require significant regulation to redirect flux when rapidly shifting from one carbon substrate to another. Many of these predictions are enzymes that are known to undergo metabolic regulation (blue; see Table S1 for identities of enzymes).

We validated RuMBA by predicting the enzymes that are regulated in *E. coli’s* canonical diauxic shift from glucose to acetate metabolism (Figs. 1 and S3). As *E. coli* grows on glucose, acetate and other fermentation products are secreted; when glucose is exhausted, the cells metabolize acetate(Holms and Bennett, 1971). When metabolism shifts from glucose to acetate metabolism, our model accurately predicted that flux diverts from isocitrate dehydrogenase to isocitrate lyase (p<<1x10^-5^; Fig. 1a-b). Furthermore, the model predicted that 131 additional proteins could be regulated to aid in the diauxic shift (Table S1), many of which have been experimentally shown to be allosterically regulated in *E. coli* (Fig. 1c; Tables S2-S3).

### PTMs occur on key enzymes in *E. coli* metabolism that require regulation

We wondered if PTMs could also regulate RuMBA-regulated enzymes. Analyzing lists of *E. coli* peptides with S/T/Y phosphorylation(Macek et al., 2008), lysine acetylation(Yu et al., 2008; Zhang et al., 2009), or lysine succinylation(Zhang et al., 2011), we found 56% of the modified proteins are metabolic (Fig. 2a), far more than expected, even when controlling for expression level (hypergeometric p<3x10^-8^). Therefore, PTMs could be important in the global regulation of metabolism. However, we note that currently, very few metabolic enzymes are known to be regulated by phosphorylation or acetylation in *E. coli* (Pisithkul et al., 2015).

**Fig. 2.**
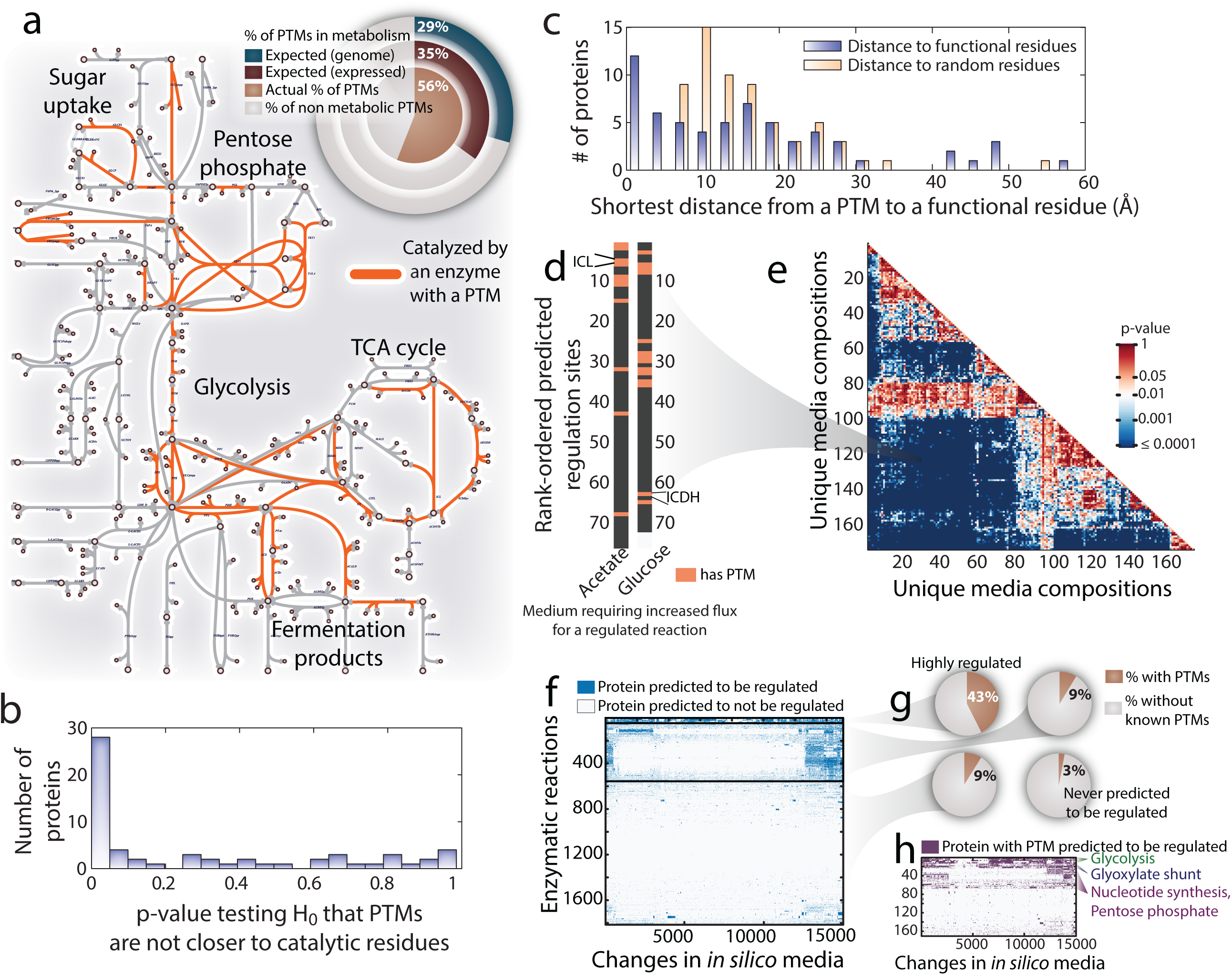
PTMs are associated with enzymes predicted to require metabolic regulation. **(a)** PTMs are found on many enzymes in central metabolism and other pathways. 56% of proteins with PTMs are in *E. coli* metabolism, which is ∼2X expected (29%) when considering protein-coding genes (or 35% when considering only expressed genes). **(b)** Across 62 metabolic protein structures, PTMs were significantly closer to functional residues (i.e., catalytic or substrate binding residues) than expected >48% of the proteins. **(c)** 37% of the proteins had PTMs closer than 10Å to functional residues. **(d)** The RuMBA algorithm provides a rank-ordered enzyme list of enzymes that should be regulated to redirect flux when rapidly shifting between nutrients. PTMs (salmon) are enriched in RuMBA predictions for the glucose-acetate diauxie, and for **(e)** 92% of the pairwise shifts between 174 media conditions. **(f)** We clustered all enzymes predicted to require regulation for at least one of the 15,051 media shifts. **(g)** 43% of the enzymes in the highly-regulated cluster have one or more PTM, while ∼9% in the less regulated clusters have PTMs. Only 3% of the enzymes that never require regulation have PTMs. **(h)** Clusters of RuMBA regulated enzymes reveal key pathway features underlying different conditions that may stimulate regulation.

Further evidence supports the functional relevance of PTMs in metabolism. First, PTM residues on RuMBA-regulated proteins are highly conserved across 1057 prokaryotic genomes (compared to non-modified S/T/Y/K residues in the same proteins, p = 0.0041; rank-sum test). Second, an analysis of PTMs on 62 proteins with available crystallographic structures (Table S3), showed many PTMs were within 10Å of catalytic site residues (Fig. 2b-c). Furthermore, at least half of the 62 metabolic proteins have PTMs that potentially disrupt or create salt bridging interactions. Third, the deletion of kinases and acetyltransferases significantly changes *E. coli* growth on individual carbon sources (Fig. S4), with some mutants increasing fitness in glucose M9 media, while others preferred M9 media with poor carbon sources, thus suggesting they may help regulate enzymes involved in metabolism in those conditions. Together, our findings suggest that (i) metabolic enzymes are enriched for PTMs, (ii) the PTMs are located where they could directly modulate catalysis or complex assembly, and (iii) kinase or acetyltransferase mutants show different phenotypes on media with specific metabolic needs.

We further investigated if PTMs could regulate metabolic flux under diverse changes in environmental conditions, beyond the glucose-acetate diauxie. We simulated changes in metabolism for 15,051 shifts between pairs of media (see Table S4) and predicted necessary regulation in each shift. PTMs are enriched among regulated enzymes in the glucose-acetate diauxie (Fig. 2d), and 92% of the other 15,050 shifts in media (hypergeometric test; FDR<0.01; Fig. 2e). Furthermore, we found that enzymes with PTMs required regulation in more conditions than enzymes without PTMs (Wilcoxon p=6x10^-6^). Indeed, enzymes cluster into four groups based on their frequency of regulation (Fig. 2f; Fig. S5-S6). In the most highly regulated cluster, 43% of the enzymes had PTMs (Fig. 2g), which is enriched in glycolytic enzymes and the glyoxylate shunt (Fig. 2h). In contrast, far fewer enzymes have experimentally measured PTM sites in clusters with enzymes requiring minimal or occasional regulation (9%) or enzymes not requiring regulation (3%). In summary, PTMs are positioned in pathways to regulate metabolism and rapidly transition between metabolic states when the nutritional environment changes.

### Protein modifications influence *in vivo* fitness in dynamic environments

We experimentally assessed when the PTMs influence cellular fitness *in vivo* by adapting MAGE(Wang et al., 2009) to perturb the PTM states of metabolic proteins. We introduced 268 targeted genetic modifications to change 134 known PTM sites in 61 proteins (Tables S5-S6) to amino acids (Fig. 3a) that (i) mimic a PTM (i.e., replacing S/T/Y residues with glutamate to mimic phosphorylation or asparagine for acetylation, referred to as “PTM-mimic”), or (ii) remove the propensity for PTM addition (i.e. replacing S/T/Y with asparagine or lysine with arginine; referred to as “PTM-null”). These targeted mutations frequently impacted organism fitness under specific media conditions, as exemplified by the PTM-null mutation at K54 of serine hydroxymethyltransferase (glyA), generated by MAGE. This mutation doubled growth rate on acetate M9 minimal medium, while not having a considerable impact on growth on glucose M9 minimal medium (Fig. 3b).

**Fig. 3.**
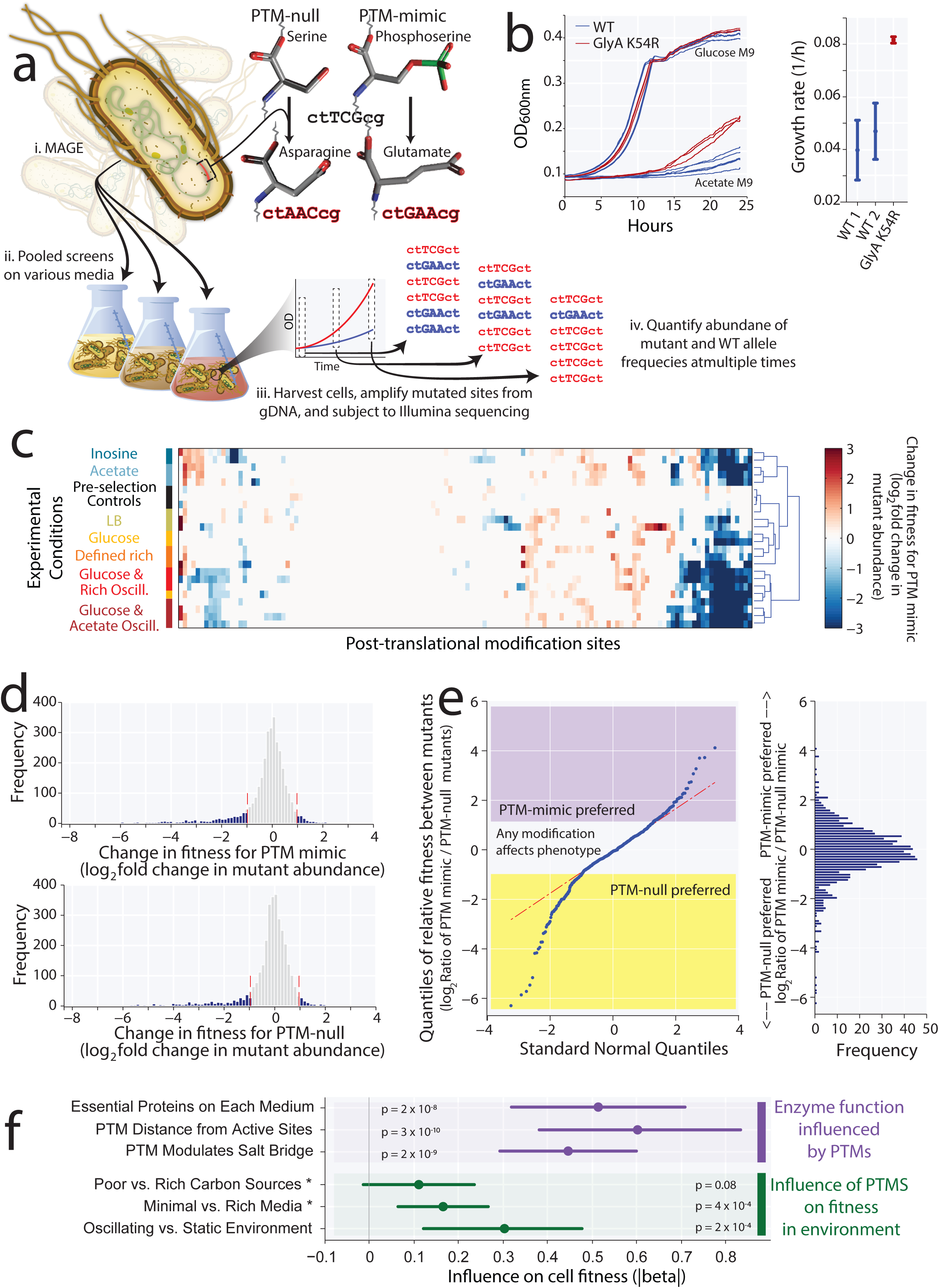
Enforcing PTM states with MAGE demonstrates PTMs are functionally relevant and impact fitness in oscillating environments. **(a)** The MAGE-generated mutant population had codons edited at PTM sites to mimic the PTM or disallow the PTM (“PTM-null”). A mutant pool harboring 268 different mutations to 134 amino acids on 61 genes was obtained and screened on diverse media conditions. Mutant abundance was quantified by amplicon-sequencing MAGE sites. **(b)** Individual mutants showed media-specific responses (e.g., enhanced growth on acetate M9 media for the GlyA K54R mutant). **(c)** The mutant pool was screened on 5 media conditions and two oscillating conditions to identify PTM sites that, when forced in a PTM-mimic or PTM-null state, had altered cellular fitness in specific conditions. **(d)** Many combinations of PTM mutations/media conditions showed >2 fold changes in fitness, and 88% of PTM sites showed a phenotype in at least one condition. **(e)** PTM sites on 35 proteins exhibited a significant preference for the PTM-mimic or PTM-null states. **(f)** A GEE analysis quantified factors influencing cell fitness after MAGE inhibited the change of PTM state. MAGE mutations only moderately impacted fitness on complex or minimal media. In comparison, the impact on fitness was almost three times as great on oscillating media when the PTM state was frozen in the PTM-mimic or PTM-null state. The localization of PTMs near active site residues on essential proteins significantly influenced fitness, especially when predicted to modulate salt bridges. Thus, the PTMs are poised to regulate the key proteins to immediately adapt to changes in growth condition.

To allow for rapid profiling of many PTM sites across several growth conditions at multiple time points, we used pooled screens (Fig. 3a) to test cell fitness effects for each of the 268 genetic changes across different media conditions. To quantify the influence of each genotype on population fitness, we measured the frequency of wild-type (WT) cells, “PTM-mimic” mutants, and “PTM-null” mutants before the screens and at 2-4 timepoints during the screens. In the screen, the fitness of many MAGE mutants was significantly impacted in specific media conditions (Fig. 3c), exhibiting positive or negative fitness in different conditions (i.e., >2 standard deviations, corresponding to 1.94 and 1.97-fold difference in abundance of mutant alleles in the “PTM-null” and “PTM-mimic” populations. respectively; Fig. 3d). Indeed, 88% of the sites showed a significant change in abundance in at least one screen. Furthermore, 35 genes had PTM sites showing significantly different impacts between “PTM-mimic” and “PTM-null” variants in at least one condition (FDR<0.05; Fig. 3e), thus suggesting that many PTM states are preferred in a specific nutrient conditions.

#### PTMs impact cell fitness by regulating key proteins in dynamic environments

What properties determine whether a PTM influences cellular fitness? To address this, we performed a global analysis of the MAGE screen data using a generalized estimating equation (GEE)(Ratcliffe and Shults, 2008) to identify how modifications impact fitness through several protein and environmental factors. The GEE is a semiparametric regression technique and allows one to control for correlation across samples with multiple time points. First, the GEE showed that modifications impacting cellular fitness are often located on proteins that are essential for growth in the medium tested (Fig. 3f). That is, enzymes critical for growth are prime targets for PTM regulation to gain a fitness advantage. Second, MAGE modifications impacting fitness often occur at structurally-relevant positions in proteins (e.g., salt bridge residues or near active site residues), which could impact growth by controlling enzyme activity. Third, modifications can influence the cell’s ability to adapt to environmental shifts. Specifically, the effects of modifications were tested within two types of nutrient environments, static (a single medium) and oscillating (media conditions were changed periodically, such as between glucose and acetate). The GEE demonstrated that when MAGE mutations force proteins to remain in a single PTM state (thereby preventing transient control at these sites), they more significantly impact cellular fitness in oscillating environments compared to static environments (Fig. 3f).

Altogether, the genetic screens show that PTMs are functionally relevant and that they regulate specific enzymes *in vivo*. Specifically, the GEE analysis of the MAGE data showed that PTMs are best positioned on enzymes in the metabolic network to control cell physiology under the growth conditions tested. Perhaps most importantly the modifications impact *in vivo* cellular fitness most when the primary nutrients change, consistent with the computational predictions of PTMs localizing to model-predicted branch points to facilitate pathway switching when the nutritional environment changes. This reinforces the notion that PTMs are required by the cell to quickly adapt to rapidly changing nutrient conditions.

### PTMs alter the molecular properties of proteins to modulate their activities

Our findings from metabolic modeling and MAGE demonstrated that many PTMs likely regulate specific enzymes to adapt to changes in nutrient availability, and the results from the analyses identify which proteins are most likely to be regulated by PTMs (Tables S3 and S7). To unravel the specific mechanisms by which the PTMs regulate the proteins, we further studied three *E. coli* proteins, for which such knowledge had never been previously unraveled. Furthermore, we highlight how similar mechanisms are likely used by the remaining proteins in our study. Specifically, we show that the PTMs (i) modulate interactions at protein interfaces, (ii) manipulate binding site conformations, or (iii) control catalytic residues. To do this, we subjected each to classical molecular dynamics simulations and *in vitro* biochemical assays for these three example proteins.

#### PTMs modulate protein interactions at dimer interfaces

Serine hydroxymethyltransferase (SHMT) requires regulation in 12% of all *in silico* substrate shifts based on RuMBA. The protein is acetylated at K54, K250, and K354, near the dimer interface, which forms part of the substrate binding domain. Using the SHMT crystallographic structure (PDB 1DFO), we analyzed long timescale molecular dynamics trajectories to analyze each acetylation site in both substrate-bound and substrate-free complexes.

K250 and K354 acetylation (where two SHMT subunits interact) affects the structure of the N-terminal domain of monomeric SHMT with an average root mean squared deviation (RMSD) of 3.5Å relative to the crystallographic structure (100ns simulation; Fig. S7). Acetylation of K250 also interferes with cofactor binding, as this residue interacts with tetrahydrofolate (THF) in 95% of the 100ns trajectory (Fig. 4a,top). Acetylation of K54 disrupts a salt bridge with E36 of the neighboring subunit, increasing the intermolecular interaction by 2-4Å, relative to WT protein (Fig. 4a and S8). *In vitro*, we found that mimicking K54 acetylation to interrupt the K54-E36 interaction reduced enzymatic activity (Fig. 4b), and the *in vivo* K54 PTM-mimic mutant leads to a significant decrease in fitness (Fig. 4c). Similarly, the MAGE mutants of K250 also decreases organism fitness *in vivo*. Thus, the molecular dynamics, *in vitro* enzymatic assays and MAGE all demonstrate that acetylation at the dimer interface of SHMT influences enzyme activity, and preventing this transient control mechanism influences cellular fitness.

**Fig. 4.**
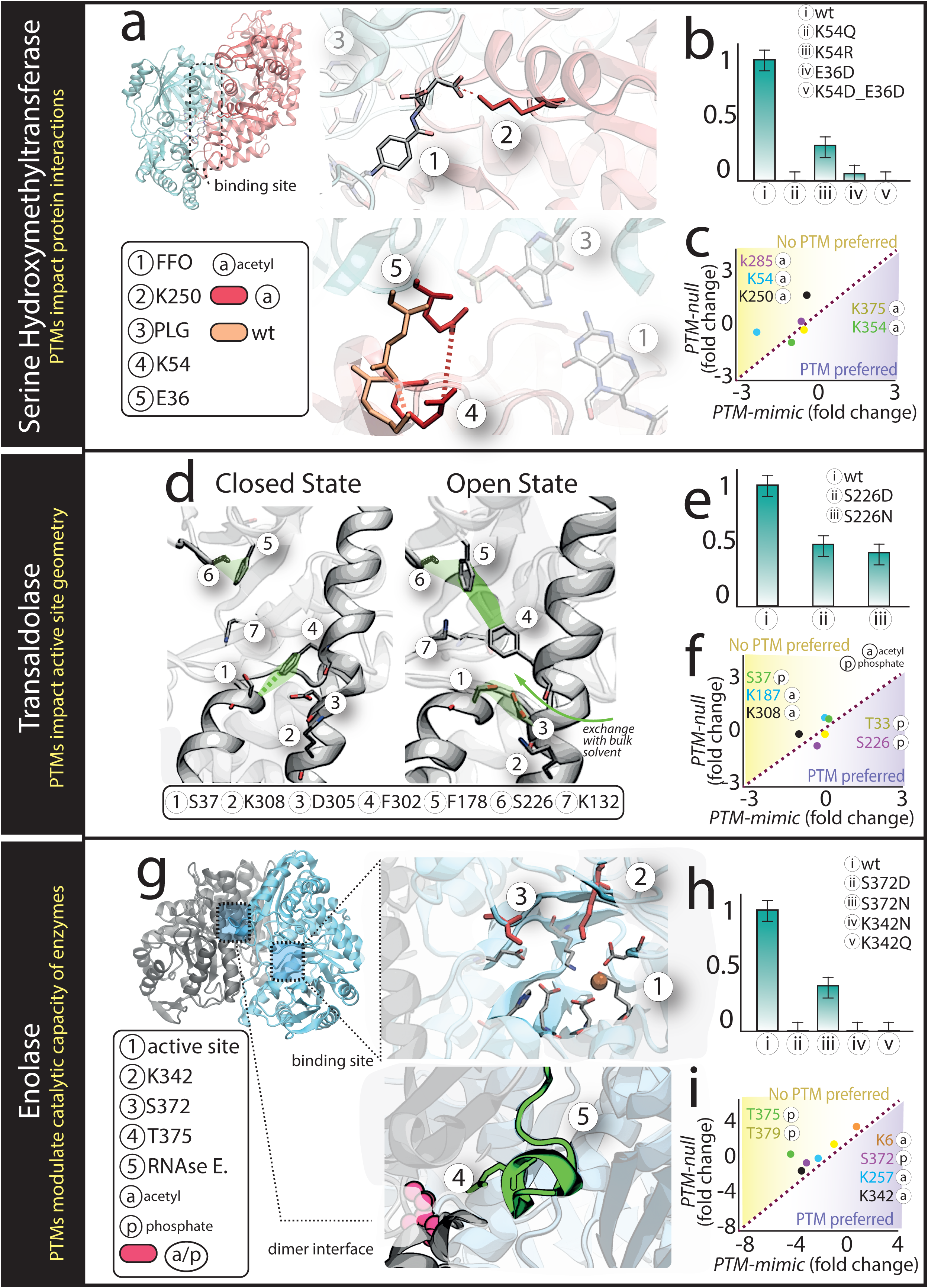
PTMs modulate protein interactions, binging site geometry or enzyme catalytic capacity. **(a)** Acetylation of K250 of serine hydroxymethyl transferase (SHMT) disrupts a salt-bridge, leading to decreased THF binding (top). K54 participates in a salt bridge with E36 of the other protein subunit (bottom), and acetylation increases the distance between K54 and E36 by >4Å (Fig. S8). **(b)** The E36-K54 interaction was disrupted *in vitro*, and catalytic activity decreased. **(c)** Average fitness for all significantly changing conditions is plotted for the PTM-mimic (x-axis) and the PTM-null (y-axis) MAGE mutants. The K54 PTM-mimic state exhibits reduced fitness, while the K250 PTM-null state significantly enhances growth. FFO= 5-formyl tetrahydrofolate and PLG= N-glycine-[3-hydroxy-2-methyl-5-phosphono-oxymethyl-pyridin-4-yl-methane]. **(d)** Interactions between six active site residues determine the open and closed states in transaldolase. The closed state involves D305-K308 and P36-F302 interactions. In the open state, S37 interacts with D305 and K308, while F302 interacts with F178, to allow the exchange of catalytic water. **(e)** Enzymatic assays show reduced activity of S226D and S226N mutants, but **(f)** the open conformation rescues the reduced activity of the S226 MAGE PTM-mimic state. The closed enzyme from the S226 PTM-null and K308 PTM-mimic reduced fitness. **(g)** (top) The catalytic site of enolase (PDB 3h8a) and (bottom) enolase in complex with a minimal binding segment of RNase E (PDB 2yfm). **(h)** PTMs modifying two catalytic residues in enolase abolish activity, as tested with *in vitro* enzyme assay and **(i)** phenotypes from MAGE mutants.

#### PTMs modulate binding site geometry

We predicted that transaldolase is regulated in 22% of all substrate shifts. Many experimentally detected PTMs occupy a shallow channel where substrates and products pass to enter/exit the active site (i.e., T33, S37, S226, K187, and K308; PDB 1ucw). The conformational state of this channel impacts substrate binding and catalysis, mediated by key residues (primarily P36, F178 and F302) that form a network of interactions between neighboring alpha helices that ultimately influence the formation of two isogenic conformational states (“closed” versus “open,” characterized by a 4Å change in substrate channel width; Fig. S9)(Lehwess-Litzmann et al., 2011).

Using classical molecular dynamics, we compared protein conformations of WT and five modified transaldolase proteins during 1.05μs (Fig. S9). The PTMs induce several molecular events. First, phosphorylation of T33 forms a salt bridge with D184 and induces large-scale structural changes in a nearby alpha helix, occluding substrate entry. Second, phosphorylation of S226 increases the interaction distance between P36 and F302 by 4Å relative to WT protein, inducing the “open” state (Fig. 4d). Lastly, acetylation of K308 disrupts a salt bridge with D305 (upheld throughout 60% of the trajectory), and interferes with interactions between P36 and F302 by inducing hydrogen bonding between D305 and S37 (Table S8).

*In vitro*, modification of S226 reduced activity by 40% (Fig. 4e), but the MAGE “PTM-mimic” state is preferred *in vivo* compared to the “PTM-null” state (Fig. 4f), presumably to maintain the “open” state, albeit at a cost of reduced activity. In contrast, forcing acetylation at K308 significantly decreases organism fitness (Fig. 4f), possibly by disrupting an “open” state formed by the salt-bridge with D305 and the D305-S37 interaction. These findings suggest that the certain PTMs act as precise, yet transient, mechano-chemical “switches” for tuning enzyme activity by impacting active site accessibility, thereby also controlling many downstream cellular processes to increase organism fitness during specific nutrient shifts.

#### PTMs control catalytic residues

Modification of catalytic residues could directly impact protein reactivity, as we found for enolase, another branch point enzyme, which we predict to be regulated in 48% of the substrate shifts. PTMs on active site residues S372 and K341 destabilize the binding of two Mg^2+^ ions, which are required for catalysis (Fig. 4g; Fig. S10). We further performed *in vitro* enzyme assays to study the effect of modifying K341 and S372 on enzyme activity (Fig. 4h; Fig. S11). Mutating S372 to aspartate arrests activity as the phosphorylation mimic introduces a negative charge near Mg^2+^. Similarly, mutating K341 to asparagine or glutamine completely arrests catalysis. These changes also cause significant changes in organism fitness *in vivo* across certain nutrient environments (Fig. 4i). These findings suggest that the direct modulation of catalytic residues provides a precise mechanism to regulate a highly sensitive branch point, such as for enolase.

## Discussion

Recent efforts in proteomics have successfully detected many PTMs in prokaryotes, but their functions have remained unclear. Here, we developed a platform to elucidate the roles of the PTMs in modulating metabolism and other cell systems.

What purpose do PTMs serve and how do they fit into the global regulatory network? We find that metabolic enzyme PTMs influence cell fitness. Our work suggests that PTMs allow cells to rapidly respond to familiar extrinsic nutrient fluctuations and intrinsic expression noise as the nutrient environment fluctuates. The PTMs satisfy an important middle ground between small-molecule mediated regulation and transcriptional regulation; providing a stable, yet rapid solution while more costly and slower processes catch up(Chubukov et al., 2013; Dekel and Alon, 2005; Lewis et al., 2010a; Zaslaver et al., 2004). Indeed, PTMs demand less energy and allow for an immediate response, compared to transcriptional and translational control mechanisms, which can take 1-2 generations in rapidly growing cells(Zaslaver et al., 2004), and likely longer to fine-tune expression(Dekel and Alon, 2005; Lewis et al., 2010a).

Our work also clearly shows from multiple angles that PTMs can directly regulate metabolism in prokaryotes, and provides examples of *how* they regulate enzyme activities. Across most changes in nutrient availability, except when media are very similar (Fig. S6), PTMs are enriched among branch point enzymes, where metabolic flux must be diverted from one pathway to another. Furthermore, the fitness effects of preventing or mimicking PTMs were greater on the MAGE screen when cells were subjected to oscillating media conditions. Finally, structural analyses and molecular simulations further supported the functional impact of the PTMs, based on proximity to important structural features and the degree of molecular changes observed upon site modifications. Together, these lines of evidence can elucidate the exact role of PTMs, and the influence they exert at both the level of individual proteins and within complex pathways.

While we find many PTMs modulate metabolism in *E. coli*, other PTMs likely regulate activities beyond metabolism. For example, the modification of specific threonine residues in enolase may impact organism fitness by influencing the binding stability of the RNAse E mediated assembly and, thus, its recruitment via the RNA degradosome (Fig. 4g, bottom; Fig. S10)(Chandran and Luisi, 2006; Kühnel and Luisi, 2001). For such cases, comparable systems level methods may elucidate the role of PTMs in non-metabolic networks (e.g., in signaling pathways). We note, however, that there may be many non-functional PTMs that occur at low stoichiometry from non-specific chemical modifications(Weinert et al., 2013). Thus, the use of systems approaches with large genetic screens can help prioritize the study of PTMs that more likely have physiologically meaningful functions.

From a systems perspective, determining states of a network requires a complete delineation of the cell parts, pathway structure, and an understanding of how the parts and pathways interact with cell environment. Our combined computational and experimental approach provides a means to reliably navigate a set of experimentally-determined PTMs and probe promising functional roles considering specific environmental perturbations. These analyses open new vistas in systems biology, empowering the systematization of biochemistry and shaping the study of PTMs in other organisms. Perhaps most important is the demonstrated ability to understand how a modification at specific sites in individual proteins can impact biological fitness, both on the molecular and physiological levels.

## Acknowledgements

The authors acknowledge support from the Swiss National Science Foundation (p2elp2_148961), the Gordon and Betty Moore Foundation (GBMF 2550.04 Life Sciences Research Foundation postdoctoral fellowship), NIH (R01-GM057089 and R35-GM119850), the US DOE (DE-FG02-02ER63445), and the Novo Nordisk Foundation Center for Biosustainability (NNF16CC0021858 and NNF10CC1016517). The authors also acknowledge NERSC computer facilities.

## Author contributions

NEL conceived and managed the research. NEL and EB led, designed, and conducted analyses. NEL, DK, EB, JX conducted experiments. RLC, HH, JTY conducted analyses. HW, CY, BOP, and GC oversaw experiments and analyses. EB and NEL wrote the manuscript. All authors read and approved of the manuscript.

## Methods

### Post-translational modifications and data on metabolic regulation

Lists of metabolic proteins with PTMs were obtained from proteomic studies of protein acetylation, phosphorylation, and succinylation in *E. coli* (Macek et al., 2008; Yu et al., 2008; Zhang et al., 2009, 2011). All reported occurrences of non-covalent metabolite-mediated metabolic regulation were obtained from Ecocyc (Keseler et al., 2005) and are reported in Table S2.

### Regulated Metabolic Branch Analysis (RuMBA)

Metabolic regulation is a rapid means to redirect flux in a metabolic network, while transcriptional regulation and regulation of enzyme abundance are processes that act on a longer time scale. Therefore, it is expected that following a shift to a new growth condition, allosteric regulation and post-translational enzyme modification will redirect flux at important branch points. The rationale for this response is that, *in vivo*, there are regular fluctuations in the cellular microenvironment and frequent environmental changes (Mitchell et al., 2009; Savageau, 1998, 1983). It would be advantageous for the cell to have a means to rapidly regulate metabolic pathway usage using reversible mechanisms while slower and more permanent regulatory mechanisms are being activated. The relative costs and timescale of a few types of regulation are given in Fig. S1.

Two methods have been developed to predict which enzymes will require significant changes in activity level following a change in carbon substrate for shorter and longer timescales, called these RuMBA and FSS, respectively. Code, compatible with the COBRA Toolbox is provided in Supplementary Data File S1.

FSS has been used previously (Bar-Even et al., 2010; Bordbar et al., 2010; Nam et al., 2012). Another method similar to FSS has also been published, showing its conceptual accuracy(Bordel et al., 2010). A brief discussion of this method provides a conceptual basis to understand RuMBA. Constraint-based modeling, the framework upon which both RuMBA and FSS are based, uses the metabolic network topology to define a space of possible phenotypes by adding a series of known biologically-relevant governing constraints (e.g., uptake rates for media components, by-product secretion rates, growth rates, etc.)(Bordbar et al., 2014; Lewis et al., 2012). This space of possible phenotypes represents all possible combinations of metabolic steady-state pathway usage that a cell can use in the given growth conditions. Assuming the constraints are accurate, the actual steady state flux distribution (or pathway usage) should be within the *in silico* solution space (Fig. S2.a). The range and distribution of flux through each reaction within these solution spaces are dependent on the constraints, such as reaction thermodynamics, metabolite uptake rates, etc. Therefore, the space is condition-specific, i.e., the various dimensions of the space might move when the model is simulated under two different growth conditions. For example, as shown in Fig. S2.b-c, the flux may be significantly higher in the second growth condition (reaction 2), or show no significant change between the two growth conditions (reaction 1).

The predicted changes in pathway usage from FSS represent the changes that lead to the optimal pathway usage in different growth conditions. However, to achieve this optimality, the activity of numerous enzymes must be fine-tuned, and often, many proteins need to be up- or down-regulated to meet this requirement. These adjustments require significant changes in transcription and translation, which can take a generation or two for entire pathways. On a shorter time scale, when changes in enzyme level are either less efficient (e.g., protein degradation) and/or not feasible to obtain, a more reasonable adaptive response involves a temporary suppression of the activity of an enzyme to avoid sending metabolites down less efficient pathways, or to boost the activity of present enzymes that will be needed in the new growth conditions. Thus, regulation at metabolic branch-points becomes of great importance, so that metabolites can be shuttled down the most efficient pathways.

RuMBA leverages this idea to compute the shift of the solution space for short-time scale changes in metabolic pathway activity at metabolic branch points. To do this, Markov chain Monte Carlo sampling of the metabolic solution space is used to obtain a uniformly distributed assessment of feasible flux values each reaction can have at steady state. To assess each branch point metabolite in the network, all reactions that can produce or consume it are identified. For example, aconitase produces isocitrate, while isocitrate dehydrogenase and isocitrate lyase both consume it (Fig. S3.a). Flux through each branch point metabolite in the network with a connectivity less than 30 is assessed. For each sample point in the solution space (Fig. S3.b-c), all incoming fluxes are summed up, as are all outgoing fluxes. Then, for each ith reaction, the fraction of total flux through the metabolite, *v*_*met*_, that is contributed by the reaction of interest, is computed as follows:

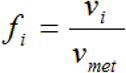

where *v*_*i*_ is the flux through reaction *i* and *f*_*i*_ is the fraction of all flux passing through the metabolite of interest, that is passing through reaction *i*. Since this is done for many random feasible sets of flux values through all of the reactions at the branch point, a distribution of *f*_*i*_ fractions is computed for each reaction for the two growth conditions of interest (Fig. S3.d). Therefore, a p-value can be computed that measures the overlap of the *f*_*i*_ values for that reaction under the given growth condition, thus quantifying how significantly the flux changes from one enzyme to another when environmental conditions change. The function of a phosphorylation event can subsequently be predicted if the change in phosphorylation is also known.

A small fraction of reactions can show miniscule, but significant changes due mostly to slight differences in predicted growth rates. Thus, the list of the regulated reactions and their associated enzymes is filtered to focus on the more significant results. Reactions that change their predicted flux level by less than 50% are filtered out from the list of reactions requiring regulation. This was done by simulating changes in reaction flux occurring in a shift between two conditions, as done previously(Bar-Even et al., 2010; Bordbar et al., 2010).

Once the flux values were normalized, the changes of fluxes between two conditions were determined as previously described (Bordbar et al., 2010). Briefly, calls on differential reaction activity were made when the distributions of feasible flux states (obtained from MCMC sampling) under two different conditions did not significantly overlap. For each metabolic reaction, a p-value was obtained by computing the probability of finding a flux value for a reaction in one condition that is equal to or more extreme than a given flux value in the second condition. Significance of p-values was adjusted for multiple hypotheses (FDR = 0.01). When the magnitude of flux changed less than 50% of the initial flux magnitude, these reactions were filtered out from the set of predicted sites of regulation and excluded from further analysis. However, results were robust for a wide range of filter levels.

To test if this method can predict the function of PTMs, three *E. coli* enzymes were identified from the literature, that undergo differential protein phosphorylation between growth on glucose and acetate. RuMBA was employed to predict the effect of phosphorylation on these three enzymes (Fig. S3.e). At late log phase, enolase has been shown to have seven times higher phosphorylation when *E. coli* was grown on glucose than when grown on acetate (Dannelly et al., 1989). *In silico*, RuMBA predicts that enolase will have a reduced flux level on acetate. Therefore, one may predict that the phosphorylation event would activate its forward flux. It was determined that when treated with acid phosphatase, enolase was inhibited(Dannelly et al., 1989). Similarly, RuMBA predicts that on acetate, the flux through isocitrate dehydrogenase (ICDHyr) decreases, while the flux through isocitrate lyase (ICL) should increase. Experimentally, the phosphorylation of ICDHyr increases and may increase for ICL (phosphorylation is high when grown on acetate, but has not been rigorously tested on glucose). Thus, it is predicted that phosphorylation of ICDHyr inhibits enzyme activity, while it activates ICL. Both of these predictions are consistent with published data (Dean et al., 1989; Hoyt and Reeves, 1988).

### Markov chain Monte Carlo sampling

The distribution of feasible fluxes for each reaction in the models used here were determined using Markov chain Monte Carlo (MCMC) sampling (Schellenberger and Palsson, 2009), as previously described (Bordbar et al., 2010; Lewis et al., 2010b), and was implemented with the COBRA Toolbox v2.0 (Schellenberger et al., 2011b). Uptake rates were used to constrain the models as detailed above. To model more realistic growth conditions (Schuster et al., 2008), suboptimal growth was modeled. Specifically, the biomass objective function (a proxy for growth rate) was provided a lower bound of 90% of the optimal growth rate as computed by flux balance analysis (Orth et al., 2010). Thus, the sampled flux distributions represented sub-optimal flux-distributions, while still modeling fluxes relevant to cell growth and maintenance.

MCMC sampling was used to simulate thousands of feasible flux distributions (referred to here as “points”) using the artificially centered hit-and-run algorithm with slight modifications, as described previously (Bordbar et al., 2010; Lewis et al., 2010b). Briefly, a set of non-uniform points was generated. Each point was subsequently moved in random directions, while remaining within the feasible flux space. To do this, a random direction is first chosen. Next, the limit for how far the point can travel in the randomly-chosen direction is calculated. Lastly, a new random point on this line is selected. This process is repeated until the set of points approaches a uniform sample of the solution space, as measured using the mixed fraction metric, which measures uniformity by measuring how many of the sample points pass through the middle line of the solution space(Schellenberger et al., 2011a). A mixed fraction of approximately 0.50 was obtained, suggesting that the space of all possible flux distributions is nearly uniformly sampled.

The distributions of sampled fluxes for each reaction were compared between two media conditions. First, flux magnitudes were normalized between each pair of media conditions (media A and B). To do this, a ratio of total flux through the metabolic network was computed and used to normalize each sample point. To compute this ratio, each sample point was taken and the magnitudes of all *n* non-loop-associated reaction fluxes were summed to acquire a value for the total network flux. For both media conditions, the median total network flux was taken and used to normalize each reaction flux for all sample points in medium *B*, as follows:

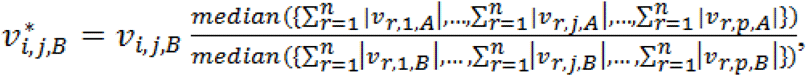

where *v*^*\**^_*i,j,B*_, is the normalized flux through reaction *i* in sample point *j* under media condition *B*, obtained after multiplying the sampled flux *v*_*i,j,B*_, by the ratio of the median total flux magnitude for the reaction for all *p* sample points under growth on medium *A* to the median total flux magnitude for the reaction for all *p* sample points under growth on medium *B*.

### Metabolic model parameterization

The genome-scale metabolic model of *E. coli* was used with published uptake and secretion rates (Feist et al., 2007). A few irreversible reactions were removed because they had reversible duplicates in the model. These include: GLCtexi, URIt2pp, URAt2pp, THMDt2pp, KAT1, INSt2pp, INDOLEt2pp, ICHORSi, CYTDt2pp, and ADNt2pp.

To identify all possible simulated media formulations in *E. coli* (Table S4), glucose uptake was set to zero in the model, and flux balance analysis was used to find which of all other carbon sources could support growth in M9 minimal media. For each of the 174 growth-supporting carbon sources, an uptake rate was set, which was consistent with uptake rate of glucose in the published model (i.e., 8 mmol gDW^-1^ hr^-1^), normalized by the number of carbons in the metabolite. For example, since glucose has 6 carbons, the uptake rate of glycerol, with 3 carbons, was set as 16 mmol gDW^-1^ hr^-1^ (which is similar to the actual reported glycerol uptake rate in M9 minimal media (Hua et al., 2006)). While this was used to standardize the media conditions, variations in carbon uptake rates did not significantly impact the results presented in this work.

### Clustering of reaction changes

An *m* x *n* matrix with m gene-reactions pairs (predicted to be regulated in at least one media shift; *m* = 1814) and *n* total media shifts (*n* = 15,051) was made, detailing in which shifts each gene-reaction pair is predicted to require regulation (FDR < 0.01). All gene-reaction pairs with at least one significantly regulated enzyme were subjected to k-means clustering (k = 3). Clustering was repeated 100 times with different seed values to find consensus clusters.

### Determination of expressed genes

For the analysis in Figure 2a, expression profiles were obtained from previous studies(Cho et al., 2009; Covert et al., 2004; Fong et al., 2005; Lewis et al., 2009). The Affymetrix CEL files were normalized using gcrma, implemented in R. Genes were considered not expressed if they did not have a mean expression level across biological replicates that were significantly higher than the five highest-expression non-*E. coli* negative control probe sets on the array (1-tail t-test; FDR = 0.05). The sets of expressed genes from each study were used to estimate the number of expressed proteins.

### Residue conservation analysis

All protein sequences of 1057 prokaryotic species were acquired from the KEGG database (Release 58.0). Homologs to all *E. coli* proteins containing at least one known PTM were identified by using the Smith-Waterman algorithm. SSEARCH35 of the FASTA suite(Pearson and Lipman, 1988) was used to determine a PID conservation for each post-translationally modified iAF1260 gene in all other genomes. The flags used in SSEARCH35 were ‘–m9 –E 1 –q –H’. When more than two proteins in one species had the same percent identity, the protein with the lowest e-value was chosen. In the rare case in which multiple proteins from a species had identical % identity scores and e-values, all qualifying proteins were included.

Each metabolic *E. coli* protein with a PTM (n=109) was then grouped with its homologs (median number of homologs for a protein = 911, 25^th^ percentile = 706, 75^th^ percentile = 1000), and the pair-wise Smith Waterman alignment between the individual *E. coli* protein and each of the homologs was used to quantify the conservation of post-translationally modified residues, as calculated (i.e., the percent of pair-wise comparisons where the aligned residue was identical in the homolog). Conservation of non-modified residues for these amino acids was calculated in an identical fashion. Relative conservation of the PTM residues on each protein was calculated by comparing their conservation to the conservation of non-PTM residues on the same protein, and a statistically significant enrichment of higher conservation was seen for PTM sites on proteins that were predicted to be regulated by RuMBA.

Conservation was done in comparison to other STYK residues on the same proteins and shown to be on average more conserved compared to other STYK on the same proteins. We have clarified this in the text.

### Salt bridge prediction and measurement of distance from PTMs to active site residues

Protein structures for modified enzymes were obtained from the Protein Data Bank. Potential salt bridges that could be disrupted by a PTM were determined by finding all residues within 4Å of a lysine or serine that could form a salt bridge. Potential new salt bridges were found by searching for basic residues within 8Å of a phosphorylated serine, threonine, or tyrosine. Distances between modified residues and all other amino acids were calculated between centroids of each amino acid. These were used to compare distance between random residues and modified residues with distances between modified residues and functional residues. Functional residues are defined as active sites on proteins, substrate binding sites, and residues which modulate enzyme activity if replaced, and were all acquired from Ecocyc, Uniprot, and the literature.

### Mutant Growth assays

Wild type *E. coli* and several mutants missing kinases, phosphatases, or acetyltransferases (Δ*aceK*, Δ*cobB*, Δ*pphA*, Δ*yeaG*, Δ*yfiQ*, Δ*yiaC*, Δ*yihE*, and Δ*ynbD*) were obtained from the Keio collection(Baba et al., 2006). Gene deletion was verified by PCR of the scar region, and strains were subsequently grown overnight M9 media, supplemented in 2g/L glucose, L-lactate, or inosine in a seeding culture. An aliquot of culture was returned to fresh media such that the OD600 was ∼0.03. Cultures were subsequently grown at 37°C with constant stirring. Turbidity was periodically measured at OD600 as a proxy for cell count, and growth rates were computed from OD measurements at mid-exponential phase.

### Strains and culture condition for MAGE

The EcNR2 strain (Wang et al., 2009) used here was a mutant of WT MG1655 *E. coli* in which the λ prophage with the bla gene was introduced via P1 transduction at the bioA/bioB gene locus and selected on ampicillin. In the strain, mutS was also replaced with a chloramphenicol resistance gene (cmR cassette). To enhance electroporation efficiency, EcNR2 was grown in LB-Lennox medium, a low salt LB-min medium with 10 g tryptone, 5 g yeast extract, 5 g NaCl, dissolved in 1 L ddH_2_O, with 50 μg/ml carbenicillin. For growth screens following MAGE, M9 minimal media was used (Teknova, catalog #M8005), supplemented with 0.1 μM biotin and carbon sources of 1.77 g/L glucose, 4 g/L NaAc*3H_2_O, or 1.58 g/L inosine. For growth selection, Azure media was also acquired from Teknova (catalog #3H5000) and supplemented with 1.77 g/L glucose. LB-Lennox was used for all LB experiments.

### Oligonucleotide design for MAGE

A panel of phosphorylation and acetylation sites were identified from previous studies(Macek et al., 2008; Yu et al., 2008; Zhang et al., 2009), and codons for the phosphorylation sites on serine and threonine or lysine acetylation were changed. Serines and threonines were changed to glutamate to mimic the phosphorylation and an asparagine to mimic the unphosphorylated residue. Lysines were converted to glutamine to mimic the acetylated state and arginine to inhibit acetylation. Codons were selected to require at least two point mutations to the gene sequence in order to ensure that subsequent sequencing of the wild-type and mutant forms would not be masked by sequencing errors. All 90-mer MAGE oligonucleotide sequences are provided for the subset of genes studied (Tables S5-S6). MAGE oligonucleotides were synthesized by Integrated DNA Technologies with standard purification. Oligos were designed to target the lagging strand and to minimize secondary structure. MAGE Oligonucleotides also contained four phosphorothioate bases at the 5’ end to enhance efficiency as described previously(Wang et al., 2009). Additional primers were designed to validate a subset of the targets using MASC-PCR (Table S9). Two sets of primers were designed to enable a two-step amplification and library preparation for amplicon sequencing and barcoding of libraries for each sample (Table S10-11). Specifically, the first set of primers were designed to amplify 99 regions containing all mutation sites targeted in our screen. At the 5’ end, each forward primer also contained the sequence 5’-CCTACACGACGCTCTTCCGATCTNNNN-3’ and each reverse primer contained the sequence 5’-GAGTTCAGACGTGTGCTCTTCCGATCT-3’. The second set of primers were designed to add the remaining sequenced needed for barcoding and next-generation sequencing.

### MAGE

MAGE was conducted as previously described (Wang et al., 2009). Specifically, cultures were initially inoculated with EcNR2 cells into 3 mL of LB-Lennox medium, and cells were grown in sterilized 10-ml polystyrene tubes at 30°C in a rotating incubator under gentle agitation until they reached an OD of 0.4 at 600nm. Cells were then heat shocked at 42°C in a shaking water bath (300 rpm) for 15 minutes. The cells were then chilled at 4°C to make them electrocompetent. One mL of cells was subsequently gently washed through several rounds of centrifugation, buffer exchanges with ice-cold ddH_2_O, and resuspension. The washed cell suspension was then mixed with 50μL single-stranded MAGE oligos (total concentration of 10μM), which were then electroporated into cells in a 1 mm gap conductive cuvette with the following setup: 1.8 kV, 200 Ω, and 25 μF. The cells were then resuspended with LB-Lennox media in preparation for further rounds of MAGE. Four rounds of MAGE were conducted. Multiplex allele-specific colony PCR (MASC-PCR) was used as previously described(Wang et al., 2012) to verify mutations and to identify specific mutants for phenotyping.

### Screen for PTM mutation fitness

We used pooled screens to assess any changes in cell fitness for each of the 268 genetic changes across multiple media conditions (e.g., LB, Azure defined rich + glucose, Glucose M9, Acetate M9, and Inosine M9) at 30°Cas well as for two oscillating conditions (Azure and glucose M9, or glucose and acetate M9). The screens were sampled at 2-4 time points (Table S12) and allele frequencies were quantified by amplifying the genes with PTM sites from the genomic DNA and sequencing the amplicons with next-generation sequencing (NGS). To obtain the final pool with all MAGE mutants, multiplexed MAGE was conducted in 5 batches, each with approximately 46 different MAGE oligos. MAGE oligos were grouped to ensure that no two oligos targeted within 100 basepairs of each other, to avoid competition between oligos in any one pool. The batches of mutants were combined and subjected to phenotypic selections.

Measurements of the allele frequency were made at three hours after electroporation and pooling and overnight storage at 4°C. Cells pellets were subsequently washed with the medium used in the screen. Cells were maintained at 30°C at exponential growth by serial dilution at regular intervals (about every three doublings; see Table S12 for values). Aliquots were saved at each dilution, and time points were selected for subsequent sequencing and analysis of allele frequencies at each PTM site.

In addition, oscillatory experiments were designed to test the fitness of the mutants when subjected to periodic changes in the nutritional environment. The oscillating conditions tested here were (i) glucose M9 and glucose-supplemented Azure chemically defined rich media and (ii) glucose M9 and acetate M9 minimal media. The experimental details are as follow. After the initial expansion of cells after the final electroporation and pooling of MAGE batches, the 24 hour time point cells were washed with the starting medium for the oscillation and allowed to grow to an OD of 0.3 at 600nm. At that point, the cell pellet was washed with the second medium and grown therein. Media were then periodically alternated after every 1-2 doublings (see Table S12 for details). More data on doubling times and the results from MAGE screen are found in Tables S12-16.

### Sequencing, alignment and quantification of variants

For each sample, cells were pelleted and DNA was isolated using the MasterPure DNA purification kit (Epicentre), and quantified using Qubit Fluorometric quantification. Sequencing libraries were prepared as follows. Genomic regions targeted by the MAGE oligos were amplified by PCR using the KAPA HiFi HotStart DNA polymerase and primers in Table S10. Amplicons were gel-quantified using ImageJ. For each sample, amplicons were pooled and a second set of PCR primers added barcodes to each sample (Table S11). Samples were gel purified, Qubit quantified, and paired-end sequenced on a HiSeq 2500.

We developed a custom DNA sequence aligner tailored to our MAGE sequencing data to map the reads to the genome and to quantify the MAGE mutants. This was done with our algorithm called KmeR-based Alignment for Multiple mismatchEs per Read (KRAMER; see code in Supplementary Data File S1). This Python-based DNA sequence aligner allows the alignment of sequencing reads with high mismatch frequency to be aligned to a predetermined set of genomic loci. The aligner takes in these loci as input and aligns Kmers derived from each sequencing read. That is, each sequencing read is broken into a set of Kmers of length k (default = 8). Of these Kmers, m (default = 8) must map to a particular locus in order for that read to be mapped. The reads can be broken into overlapping Kmers by specifying o (default = no overlap). For the results shown here, values for k and m were varied to provide the best results; k=8 and m=8 were chosen after sensitivity analysis

After assigning a locus to each sequencing read, each read is compared to the wild type locus to determine if a particular target site in the locus perfectly matches a site in the read. Specifically, in this study, we used MAGE to change at least one codon in each gene. Thus, we searched for perfect matches surrounding the site of the modified codon, and then also looked to see if the site of the modification had the WT codon (Ser,Thr, or Lys), the codon for the PTM mimic (Glu or Gln), or the codon for the amino acid that cannot be post-translationally modified (Asp or Arg). The algorithm uses a parameter called targetsize t (default = 9), which is specified to be the length of the stretch of target DNA that will be matched; in this implementation, t is an odd number from 5 to 43 (e.g., t = 5 would have the target codon with one flanking nucleotide on both ends). To aid in quality control assessment, reads that map to the *E. coli* genome but that do not map to the targeted loci are saved to a separate file, thus allowing further analysis and identification of potential contaminants. Similarly, reads that do not map to the *E. coli* genome or MAGE target sites are written to a file for quality assessment.

The implementation provided in this work allows for other optional arguments:

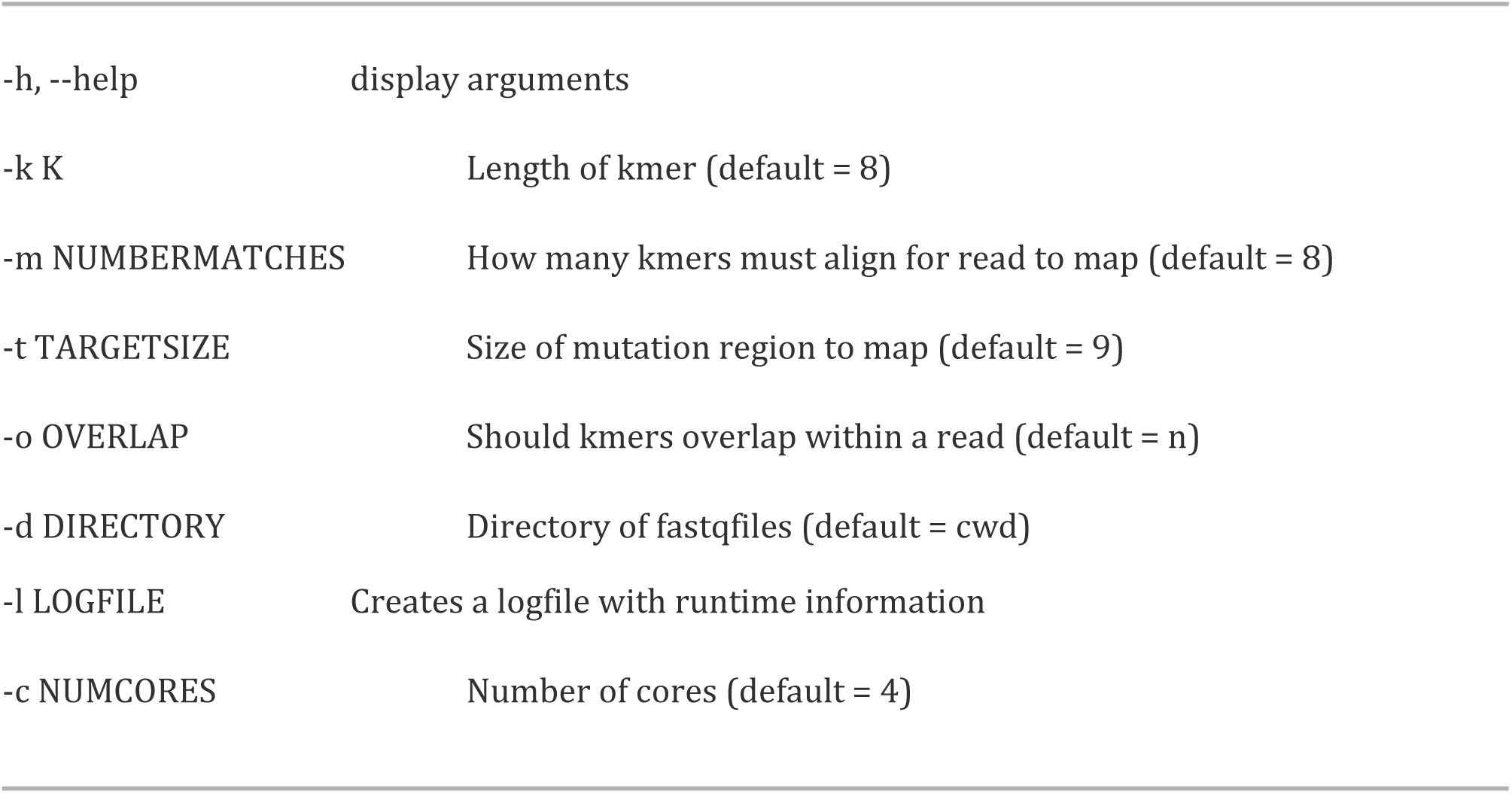

After quantifying the allele frequency for each sample, the allele frequencies were median-normalized, and fold change in frequency was determined by log transforming the allele frequencies and after subtracting the mean frequencies of the control samples (hour 3, pre and post incubation at 4°C).

### Identification of covariates modulating the impact of loss of PTM switching

We first identified several biological features for each experiment, gene, and modification site for the MAGE screens. These included the following phenotypic features for experimental samples: 1) whether the experiment was a steady growth condition or oscillating, 2) if the media included glucose or an alternative poor carbon substrate, 3) if the media was M9 minimal media or a rich medium, and 4) the number of doublings seen by the sample after the start of the time course. In addition, we considered, for each PTM, if the modification was phosphorylation or acetylation, and if the modification was on a gene that is predicted to be essential for the given growth condition, based on flux balance analysis simulations (Lewis et al., 2012).

### GEE analysis of MAGE screen data

To identify features that best explained the variation in phenotypic impacts of the MAGE mutations, the generalized estimating equation was used with Markov correlation structure using the GEEQBOX package in MATLAB(Ratcliffe and Shults, 2008). This model identified features that best explained the variation in phenotypic impacts of the MAGE mutations. This model was used to control for the multiple measurements of each experiment while controlling for variation in number of doublings across the samples.

A univariate pre-screening was conducted to assess the contribution of each experimental and biological feature. Since each sample was measured at multiple time points, the generalized estimating equation was used with the Markov correlation structure(Hanley et al., 2003) to account for correlation between time points. Biological features that were not significant in the univariate pre-screening were eliminated from further analysis. Significant variables were subsequently assessed for multicollinearity to eliminate redundant variables. Following the pre-screening, a few features were identified as providing a significant contribution to fitness of mutants in the screen. These included 1) whether the cells were grown in a single growth condition or oscillating media, 2) whether the media contained glucose or a poor carbon source, 3) whether the media was rich or minimal media, 4) if the PTMs were on essential genes for the given growth condition, 5) the proximity of the PTM to active site residues, and 6) whether the PTM is predicted to modulate salt bridges. The significant media conditions were multicollinear and two models were analyzed including only one of the two correlating features. In the final models, analyses comparing poor vs. rich carbon sources and minimal vs. complex media were correlated and therefore were analyzed in separate models.

### Molecular Dynamics Simulations

Classical molecular dynamics simulations were performed starting from the crystal structure of all proteins. The individual mutations were manually changed according to the post-translational modification of interest. Parameters for the phosphorylated amino acids were based on the parametrization of Homeyer et al. (Homeyer et al., 2006). Using PROPKA (Bas et al., 2008; Li et al., 2005; Olsson et al., 2011) we estimated that all of the residues adopt the default protonation states. All other non-standard parameters (i.e. for substrates) were calculated per procedures used for the generation of the parm99 parameters and recommended in the AMBER manual. RESP (Bayly et al., 1993; Cieplak et al., 1995; Cornell et al., 1993) charges were generated by performing a three stage RESP fit on two HF/6-31G* optimized structures. Simulations were performed for both substrate-bound and substrate-free states. Each structure was solvated with TIP3P water and, depending on the total charge of the system, either 14, 7, 20, Na^+^ ions, achieved system neutrality, (for apo serine hydroxymethyltransferase (SHMT), transaldolase and enolase, respectively), in an orthothrombic periodic box (dimensions for SHMT: 94 × 95 × 114 Å; dimensions for transaldolase: 89 x 74 x 75 Å; dimensions for enolase: 110 x 101 x 89 Å). The particle mesh Ewald (PME) method(Darden et al., 1993; Essmann et al., 1995), with a nonbonded cutoff of 12 Å, was used with periodic boundary conditions and the Langevin piston Nosé–Hoover method (Feller et al., 1995; Martyna et al., 1994; Nosé, 1984) to ensure constant pressure and temperature conditions. For each system, GPU-enabled PMEMD molecular dynamics was performed (Salomon-Ferrer et al., 2013), using the AMBER 99sb force field (Hornak et al., 2006; Wang et al., 2000) for 50-120 ns per protein state (i.e., substrate-bound versus substrate-free in wild-type or modified variant proteins).

### Enzymatic Assays

Enolase activity was assayed by measuring the conversion of 2-PGE to PEP at 25°C as described previously(Liu et al., 2012) with modifications. The reaction mixture contained 1 mM 2-PGE in reaction buffer (100 mM HEPES buffer, pH 8.5, 7.7 mM KCl, 10 mM MgSO_4_, prewarmed to 25°C), and enolase was added to initiate the reaction. The reaction was monitored spectrophotometrically by measuring absorbance at 240 nm for the production of PEP at 30 sec intervals for 10 min.

For transaldolase, the reverse reaction catalyzed by transaldolase was tested at room temperature as described previously (Huang et al., 2008) with some modifications. The reaction mixture contained 5 mM D-fructose-6-phosphate, 0.2 mM erythrose-4-phosphate, 0.1 mM NADH, and 10 μg of α-glycerolphosphate dehydrogenase-triosephosphate isomerase (Sigma) in reaction buffer (40 mM triethanolamine, pH 7.6, 5 mM EDTA), and transaldolase was added to initiate the reaction. The reaction was monitored spectrophotometrically by measuring absorbance at 340 nm at 30 sec intervals for 10 min.

Serine hydroxymethyltransferase activity of THF-dependent cleavage was measured as described previously(Schirch, 1971) with some modifications. The reaction mixture in a final volume of 75 μl consisted of 0.3 mM pyridoxal phosphate, 40 mM mercaptoethanol, 15 mM serine and serine hydroxymethyltransferase in reaction buffer (10 mm potassium phosphate, pH 7.3, 0.5 mM EDTA). After a 5-minute incubation at 37°C, 1 mM THF was added to initiate the reaction. The reaction was stopped after 2 minutes by the addition of 100 μl of pH 9.5 carbonate buffer. 20 μl of 2 mM NADP^+^ and enough methylene tetrahydrofolate dehydrogenase were then added to carry out the auxiliary reaction and the increase in absorbance at 340 nm was followed to completion.

